# Macrophage Migration Inhibitory Factor (MIF)-CD74 Signaling Pathway Mediates Trabecular Meshwork Dysfunction in Glaucoma

**DOI:** 10.64898/2026.03.18.712673

**Authors:** Monu Monu, Lal Krishan Kumar, Prince Kumar, Gulab Zode, Pawan Kumar Singh

**Affiliations:** Department of Ophthalmology, Mason Eye Institute, University of Missouri School of Medicine, Columbia, Missouri, USA; Gavin Herbert Eye Institute-Center for Translational Vision Research, Department of Ophthalmology, University of California Irvine School of Medicine, Irvine, California, USA

**Author notes:** **Address Correspondence to: Pawan Kumar Singh, Ph.D.,** Department of Ophthalmology, Mason Eye Institute, University of Missouri School of Medicine, 1 Hospital Dr, Columbia, Missouri-65212, USA. These authors contributed equally (Monu Monu, Lal Krishan Kumar, Prince Kumar).

**Keywords:** Glaucoma, intraocular pressure, trabecular meshwork, MIF, CD74, Blimp-1, agmatine, thiamine, 4-IPP, myocilin, TGF-β2, fibrotic

## Abstract

**Purpose:** Macrophage migration inhibitory factor (MIF) is a pleiotropic cytokine implicated in many inflammatory and fibrotic diseases; however, its role in primary open-angle glaucoma (POAG) and trabecular meshwork (TM) dysfunction remains unknown. In this study, we investigated whether MIF-CD74 signaling regulates TM pathobiology through modulation of the transcription factor, Blimp-1, and downstream cytoskeletal reorganization and extracellular matrix (ECM) remodeling.

**Method:** Primary human TM cells (HTMC) were exposed to glaucomatous stressors, including TGF-β2, rMIF, or a pro-inflammatory milieu. Expression of MIF, its receptor CD74, and Blimp-1 was measured by qPCR and immunoblotting. ECM proteins and phosphorylated myosin-light chain (pMLC) were evaluated by immunofluorescence staining. *In vivo*, MIF-CD74 and Blimp-1 expression were examined in the TM/anterior segment (AS) tissue of Tg.CreMYOC^Y437H^ and lentiviral (LV)-TGF-β2-induced ocular hypertension (OHT) mouse models. Functional involvement of MIF signaling in TM pathobiology was examined using the irreversible MIF inhibitor 4-IPP and the immunomodulatory metabolites agmatine and thiamine.

**Results:** Glaucomatous stressors significantly upregulated MIF and CD74 expression with concomitant suppression of Blimp-1 in HTMC. Similarly, TM/AS tissue from both OHT models (Tg.CreMYOC^Y437H^ and LV-TGF-β2) demonstrated increased MIF-CD74 expression accompanied by reduced Blimp-1 levels. Activation of MIF-CD74 signaling triggered pro-inflammatory and cell death pathways and promoted ECM remodeling, characterized by increased fibrotic protein expression and enhanced RhoA/ROCK-mediated MLC phosphorylation, indicating modulation of TM contractility. Pharmacological inhibition of MIF attenuated inflammatory signaling, reduced ECM deposition and cytoskeletal remodeling, and suppressed RhoA/ROCK/MLC activation, restoring a protective TM phenotype.

**Conclusion:** Our findings identify MIF-CD74 signaling as a previously unrecognized regulator of TM dysfunction in POAG. MIF-mediated suppression of Blimp-1 mechanistically links inflammatory signaling to cytoskeletal contractility and fibrotic ECM remodeling, key determinants of aqueous humor outflow resistance. Targeting the MIF-CD74/Blimp-1 axis may represent a novel therapeutic strategy to restore TM homeostasis and reduce intraocular pressure in glaucoma.

## Introduction

Primary open-angle glaucoma (POAG) is the most prevalent form of glaucoma and a leading cause of irreversible blindness worldwide, particularly among aging populations. Current estimates indicate that glaucoma affects ∼80 million people globally, with projections exceeding 112 million by 2040^1, 2^. Elevated intraocular pressure (IOP), resulting from trabecular meshwork (TM) dysfunction, remains the primary modifiable risk factor for POAG^3^. Although multiple risk factors, including age, race/ethnicity, family history, and pre-existing ocular conditions, contribute to disease susceptibility^4^, the molecular mechanisms underlying TM dysfunction in POAG remain poorly understood. Age-associated oxidative stress, vascular damage, and mitochondrial dysfunction are hypothesized to induce TM cell apoptosis, ECM accumulation, and cytoskeletal stiffening, collectively increasing aqueous humor outflow resistance and elevating IOP^5, 6^. In addition, various other intrinsic mechanisms, such as immune activation, ER stress, and metabolic dysregulation, have been proposed to further exacerbate TM pathology^7–10^. Despite evidence linking immune dysregulation to POAG, the specific inflammatory pathways driving TM remodeling and degeneration remain largely unknown. Moreover, current glaucoma therapies are primarily limited to lowering IOP by either reducing aqueous humor production or enhancing outflow, which can slow disease progression but do not address the underlying TM pathology. Because TM dysfunction drives outflow resistance and contributes directly to glaucoma progression, understanding the immune and inflammatory mechanisms that disrupt TM homeostasis is critical. Elucidating these pathways is essential for developing novel, disease-modifying therapies that go beyond IOP reduction to restore TM function, an unmet clinical need in glaucoma management.

Macrophage migration inhibitory factor (MIF) is a pleiotropic pro-inflammatory cytokine expressed by multiple immune cells, including monocytes, macrophages, T cells, B cells, and neutrophils^11, 12^. In ocular tissue, MIF is expressed by Müller glia, astrocytes, retinal pigmented epithelium, and TM cells^13, 14^. MIF signals through canonical receptors, CD74, and CD44, as well as non-canonical receptors CXCR2, CXCR4, CXCR7, and can also act via receptor-independent endocytosis^15, 16^. Upon activation, MIF activates MAP kinases (ERK1/2, p38, JNK) and PI3K pathways, resulting in nuclear translocation of transcription factors such as NFκB, STAT3, and AKT, and upregulation of pro-inflammatory mediators including TNFα, IL-2, IL-6, IL-8, IFN-γ, IL-17, and nitric oxide^17, 18^. MIF further potentiates inflammation through TLR4 activation, NLRP3 inflammasome engagement and IL-1β production, and suppression of anti-inflammatory cytokine IL-10^18^ ^19^. Dysregulated MIF signaling has been implicated in the pathogenesis of autoimmune, inflammatory, and neurodegenerative diseases, as well as a few ocular pathologies ^18, 20, 21^; however, its role in TM dysfunction and POAG pathogenesis remains uninvestigated.

B lymphocyte-induced maturation protein-1 (Blimp-1), encoded by PRDM1, is a transcription regulator involved in immune modulation, cellular differentiation, and stress responses^22, 23^. Blimp-1 contributes to the regulation of IL-10 expression and other anti-inflammatory pathways^22, 23^. Although MIF is known to suppress IL-10, whether MIF signaling directly modulates Blimp-1 has not been examined in any disease context. Our preliminary spatial transcriptomic analysis of glaucomatous human TM tissue and glycoproteomic profiling of primary HTMCs exposed to glaucomatous stress revealed altered MIF expression and glycosylation patterns, implicating MIF as a potential mediator of TM pathology^24^ (unpublished pilot data). These observations prompted us to investigate the potential role of the MIF-CD74/Blimp-1 axis in TM pathobiology.

In the present study, we examined the role of MIF-CD74 signaling in TM pathogenesis using primary HTMC and two complementary *in vivo* OHT models, Tg.CreMYOC^Y437H^ and lentiviral TGF-β2 induced models. Our findings demonstrate that glaucomatous stressors activate MIF-CD74 signaling, leading to suppression of the protective transcription factor, Blimp-1, promoting TM inflammation, activation of cell death pathways, increased ECM deposition, and enhanced RhoA/ROCK-mediated myosin light chain phosphorylation, indicating altered TM stiffness and contractility. Pharmacologic inhibition of MIF using 4-IPP or immunomodulation with agmatine or thiamine significantly attenuated these pathogenic responses, reducing TM inflammation and fibrotic remodeling. Collectively, our findings identify the MIF-CD74/Blimp-1 axis as a previously unrecognized regulator of TM dysfunction and suggest that targeting MIF signaling may represent a promising therapeutic strategy to restore TM homeostasis and prevent glaucoma progression.

## Results

### Ocular hypertension activates MIF-CD74 signaling and suppresses Blimp-1 in mouse TM

To determine whether glaucomatous stress activates MIF-CD74 signaling *in vivo*, we utilized two complementary mouse models of ocular hypertension (OHT): Tg.CreMYOC^Y437H^ and lentiviral TGF-β2-indcued OHT. The Tg.CreMYOC^Y437H^ model was generated using a TARGATT site-specific knock-in strategy (**Fig. 1A**), as reported previously^25^. In this model, a single intravitreal injection of HAd5 expressing Cre recombinase induces mutant myocilin expression in the TM of Tg.CreMYOC^Y437H^ mice. Successful induction of mutant MYOC was confirmed by MYOC-DsRed fluorescent protein expression in Ad5-Cre-injected mice compared with Ad5-empty vector controls (**Fig. 1B, C**). Longitudinal IOP measurements demonstrated significant and sustained IOP elevation in Tg.CreMYOC^Y437H^ mice relative to controls (**Fig. 1D**). Following IOP elevation, we evaluated MIF-CD74 and Blimp-1 expression in TM/anterior segment (AS) tissues of these mice. The immunofluorescence analysis revealed increased MIF expression in the TM, with co-localization of MIF and MYOC-DsRed in Tg.CreMYOC^Y437H^ mice (**Fig. 1E**), indicating activation of MIF under glaucomatous conditions. To further evaluate MIF-CD74/Blimp-1 signaling, we quantified transcript and protein expression levels in AS tissue by qPCR and immunoblotting, respectively. Our results demonstrate that Tg.CreMYOC^Y437H^ mice exhibit significant upregulation of MIF and CD74, accompanied by concurrent suppression of Blimp-1 at both the mRNA (**Fig. 1F**) and protein levels (**Fig. 1G, H**) compared with empty vector controls.

**Fig. 1:**
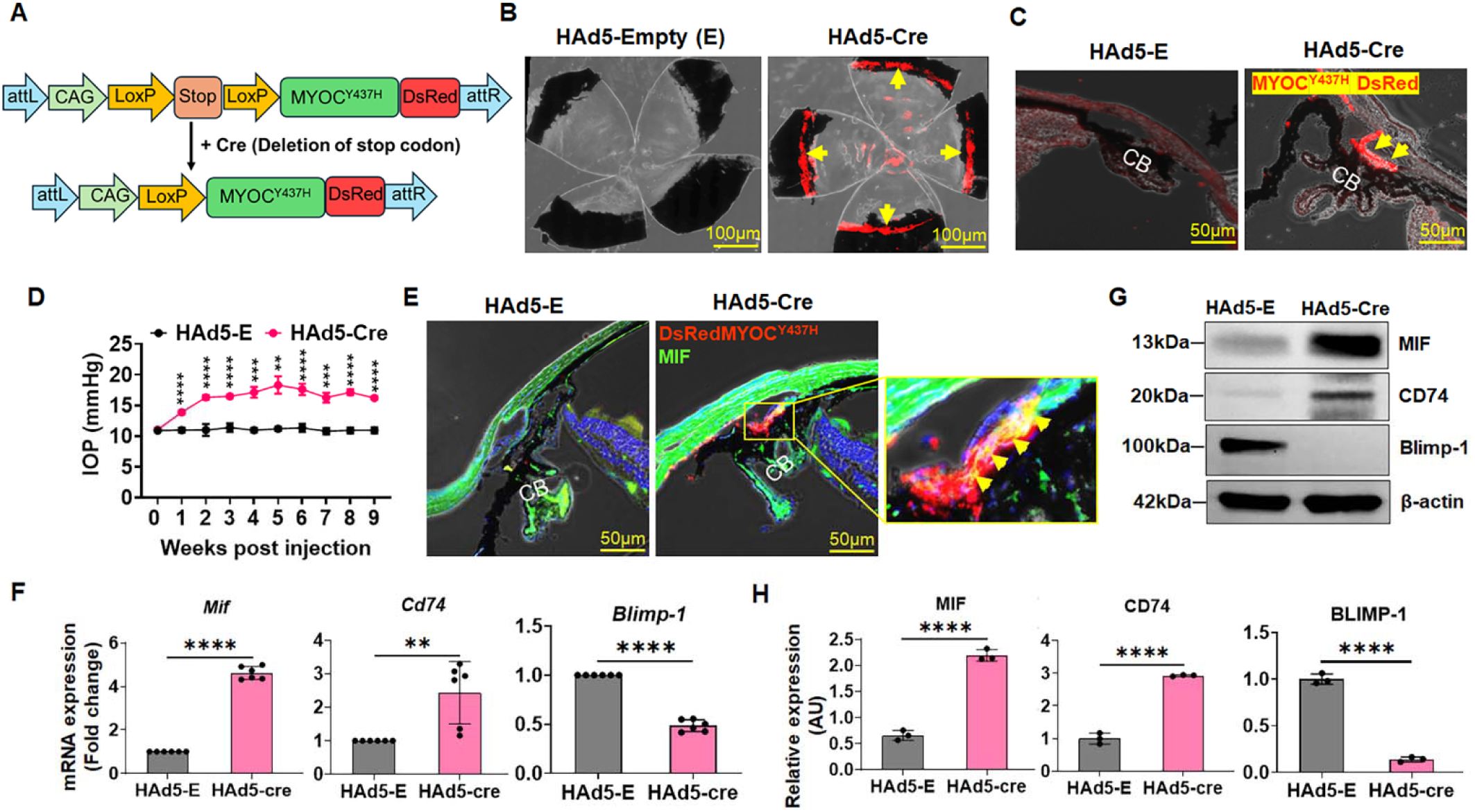
MIF-CD74/Blimp-1 axis is dysregulated in Tg.CreMYOC^Y437H^ mouse model of OHT. The Tg.CreMYOC^Y437H^ mice (n=12/group) were injected intravitreally with either HAd5-Cre or HAd5-empty vectors. (**A**) Schematic of the knock-in cassette showing DsRed-tagged human Y437H-mutant MYOC inserted at the transcriptionally active H11 locus. A transcriptional stop cassette prevents expression of the MYOC-DsRed fusion protein until Cre recombinase excises the stop sequence, enabling TM-specific expression of mutant MYOC. (**B-C**) Fluorescence imaging of whole-mount anterior segments (**B**) and ocular cross-sections (**C**) demonstrates robust DsRed-MYOC^Y437H^ expression in Tg.CreMYOC^Y437H^ mice (indicated with yellow arrows), compared with Ad5-empty controls. Scale bars: 100 μm (B), 50 μm (C). (**D**) Weekly IOP measurements show significant and sustained IOP elevation in Cre-injected Tg.CreMYOC^Y437H^ mice compared with empty vector controls. (**E**) Immunofluorescence staining of anterior segment cross-sections shows MIF expression co-localizing with DsRed-MYOC^Y437H^ in the TM of Tg.CreMYOC^Y437H^ mice. (**F-H**) qPCR (**F**) and western blot (**G, H**) analyses of anterior segment tissues demonstrate upregulation of MIF and CD74 with concurrent suppression of Blimp-1 in Tg.CreMYOC^Y437H^ mice. Western blot bands were quantified by densitometric analysis using ImageJ and normalized to β-actin, and data are presented as relative protein expression (AU) (**H**). Data represent mean ± SD. **p<0.005, and ****p<0.0001. Two-way ANOVA with Tukey’s multiple-comparison test (D), unpaired t-test (G, H).

To validate these findings in an independent OHT model, we employed the lentiviral TGF-β2 OHT model^26^. To induce OHT, wild-type C57BL/6 mice received a single intravitreal injection of LV-CMV-hTGF-β2^C226/228S^ (LV-TGF-β2), while LV-CMV-Null (LV-null) injected mice served as controls. As expected, LV-TGF-β2 injection induced significant and sustained IOP elevation compared with LV-null mice (**Fig. 2A**). To assess the expression of MIF in this OHT model, we performed co-immunostaining using anti-MIF antibody with TM marker α-SMA. Our immunofluorescence staining of AS tissue revealed increased MIF expression in the TM of LV-TGF-β2-injected mice, demonstrated by co-localization of MIF with α-SMA (**Fig. 2B**). Consistent with the Tg.CreMYOC^Y437H^ model, our qPCR and immunoblot analyses showed significant induction of MIF and CD74 with concomitant suppression of Blimp-1 at both transcript (**Fig. 2C**) and protein levels (**Fig. 2D, E**) in LV-TGF-β2-treated mice. Given the well-established profibrotic role of TGF-β2 in glaucoma, we further assessed fibrotic and ECM-associated markers in AS tissues. Our results show that LV-TGF-β2-treated mice exhibit significantly increased expression of α-SMA, MYOC, and fibronectin-1 at both the mRNA and protein levels (**Fig. 2C-E**), confirming induction of fibrotic remodeling in parallel with activation of MIF-CD74 signaling in this model.

**Fig. 2:**
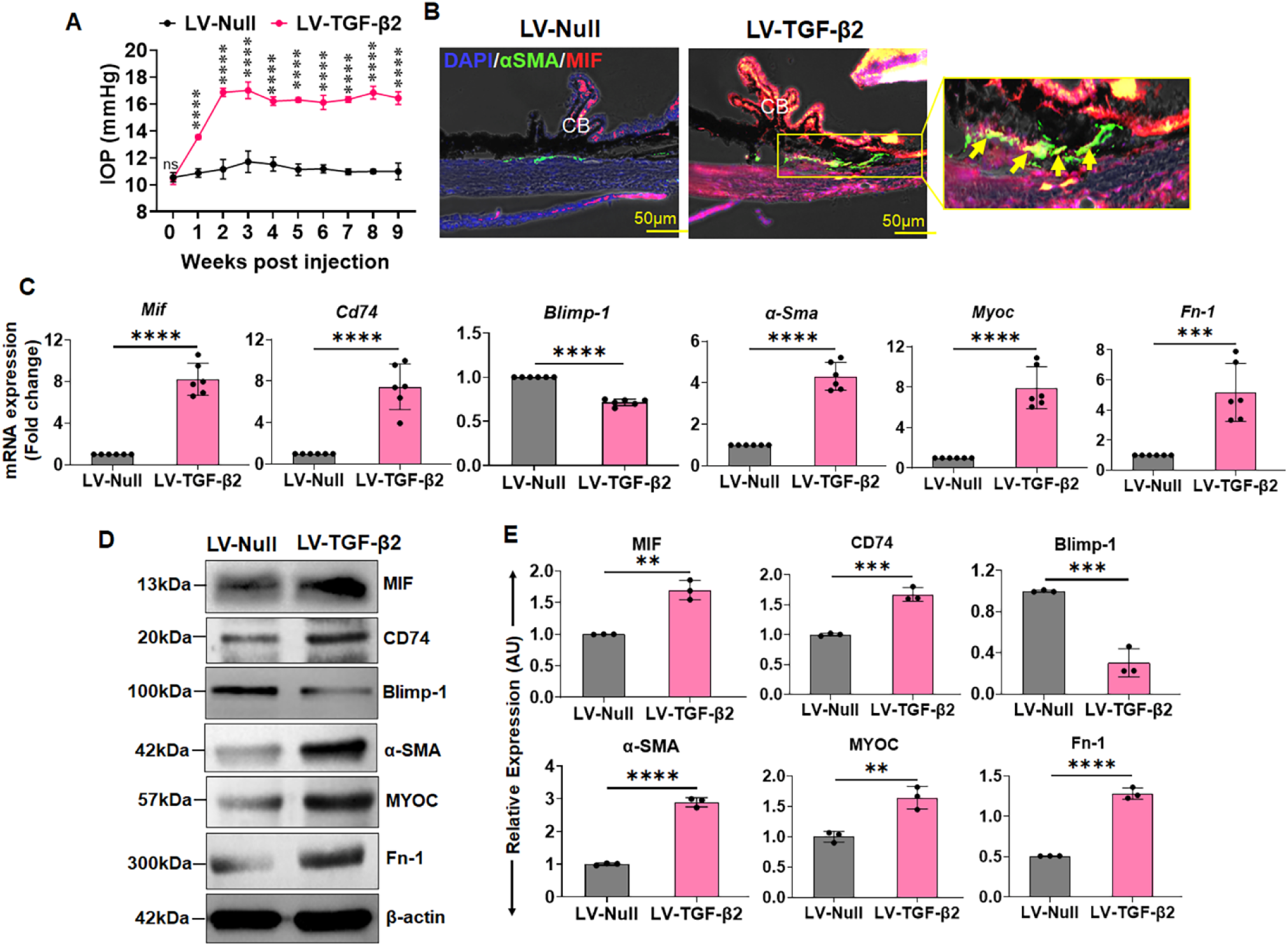
MIF-CD74/Blimp-1 axis is dysregulated in lentiviral-TGF-β2-induced mouse model of OHT. Wild-type C57BL/6 mice (n=12/group) were injected intravitreally with either LV-TGF-β2 or LV-null vectors. (**A**) Weekly IOP measurements demonstrate significant and sustained IOP elevation in LV-TGF-β2-injected mice compared with LV-null controls. (**B**) Immunofluorescence staining of anterior segment cross-sections for α-SMA and MIF shows co-localization of MIF with the TM marker α-SMA, indicating increased MIF expression in the TM of LV-TGF-β2 injected mice. (**C-E**) qPCR (**C**) and western blot (**D, E**) analyses of anterior segment tissues show upregulation of MIF and CD74 accompanied by suppression of Blimp-1 in LV-TGF-β2-injected mice. Increased expression of profibrotic markers, including α-SMA, Fn-1, and MYOC, was also evident in LV-TGF-β2-injected mice. Western blot bands were quantified by densitometric analysis using ImageJ and normalized to the housekeeping control β-actin, and protein levels are presented as relative expression (AU) (**E**). Data represent mean ± SD. **p<0.005, ***p<0.0005, and ****p<0.0001. Two-way ANOVA with Tukey’s multiple-comparison test (A), unpaired t-test (C, E).

Collectively, these findings demonstrate that experimental OHT induces pathological activation of MIF-CD74 signaling with concurrent suppression of Blimp-1, establishing this axis as a key molecular feature of glaucomatous TM pathology *in vivo*.

### Glaucomatous stressors activate MIF-CD74 signaling and suppress Blimp-1 in primary HTMCs

To further examine the impact of glaucomatous stress on MIF-CD74 signaling *in vitro*, we challenged the primary HTMCs with various pathologic stressors, including TGF-β2 (10 ng/mL, 24h), a pro-inflammatory cytokine cocktail (TNF-α, IL-1β, IL-6, IL-17A; 100 ng/mL, 24h), or recombinant MIF (rMIF, 100 ng/mL, 24h). Following treatment, the expression of MIF, CD74, and Blimp-1 was assessed by qPCR and immunoblotting. Our results revealed that exposure to glaucomatous stressors resulted in robust upregulation of MIF and CD74, with concomitant suppression of Blimp-1 at both the transcript (**Fig. 3A**) and protein (**Fig. 3B**) levels. We further confirmed MIF expression by immunofluorescence staining. Our data showed increased MIF expression in HTMCs following TGF-β2 or the inflammatory cytokine exposure (**Fig. 3C**). These results indicate that, consistent with our *in vivo* OHT models, glaucomatous insults activate MIF-CD74 signaling while concurrently suppressing the protective transcription factor Blimp-1 in TM cells.

**Fig. 3:**
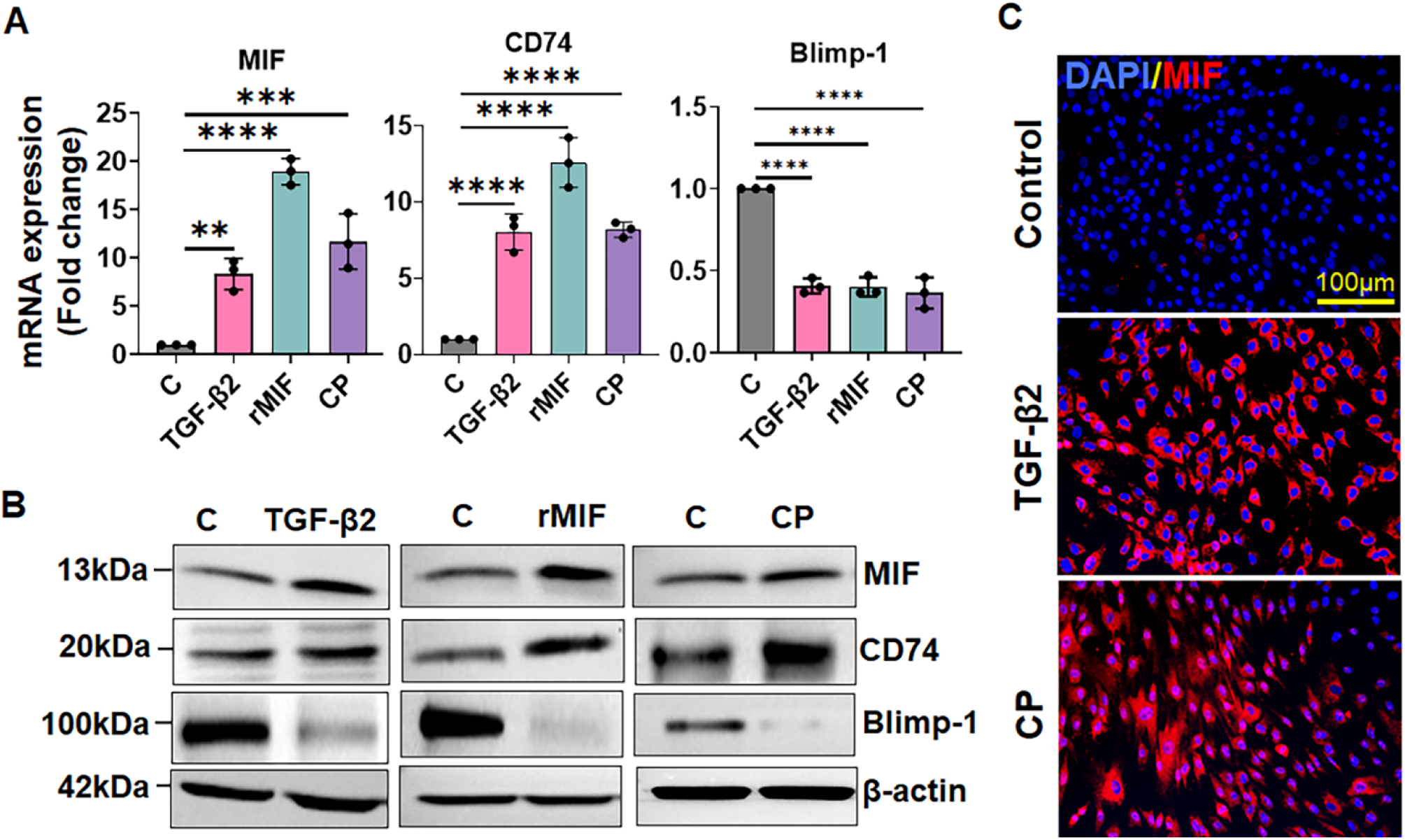
Glaucomatous stressors induce MIF-CD74 expression and suppress Blimp-1 in HTMCs. Primary HTMCs (n=3) were challenged with TGF-β2 (10ng/mL), rMIF (100ng/mL), or pro-inflammatory cytokine pool (CP; TNF-α, IL-1β, IL-6, and IL-17 100ng/mL) for 24h. **(A)** qPCR analysis shows significant upregulation of MIF and CD74 transcripts accompanied by reduced Blimp-1 expression in response to glaucomatous stressors. (**B**) Immunofluorescence staining demonstrates increased MIF expression following TGF-β2 or CP treatment. **(C)** Western blot analysis confirms induction of MIF and CD74 with concurrent suppression of Blimp-1 at the protein level. Data represent mean ± SD. **p<0.005, ***p<0.0005, and ***p<0.0001, One-way ANOVA with Dunnett’s multiple-comparison test.

### MIF promotes, while MIF inhibition attenuates, ECM deposition and cytoskeletal remodeling in HTMCs

ECM accumulation and cytoskeletal remodeling are hallmarks of TM dysfunction, which contribute to increased tissue stiffness, reduced cellular contractility, and elevated AH outflow resistance, ultimately increasing IOP. Given our observation of MIF-CD74 activation in TM, we next examined whether MIF directly promotes TM profibrotic responses. To assess this, we challenged primary HTMCs with rMIF in the presence or absence of the irreversible MIF-specific inhibitor 4-IPP, as well as two immunomodulatory metabolites, agmatine and thiamine, that we recently identified in our glaucoma patient metabolomics studies^27^. 4-IPP covalently modifies MIF, preventing CD74 binding and permanently inactivating its catalytic biological functions, whereas the effects of agmatine and thiamin on MIF activity remain unknown. Following treatment, we assessed the fibrotic and ECM response via immunofluorescence staining and qPCR. Our results demonstrate that exposure to rMIF significantly upregulated fibrotic markers, as evidenced by a marked increase in α-SMA expression (**Fig. 4A, B**). In contrast, pharmacological inhibition of MIF by 4-IPP or treatment with agmatine or thiamine drastically reduced α-SMA expression. Immunofluorescence staining further demonstrated that rMIF induced ECM and cytoskeletal proteins, including fibronectin and F-actin, whereas these effects were suppressed by MIF inhibition (**Fig. 4A, B**). Consistent with these observations, qPCR analysis showed significant induction of fibrotic and ECM transcripts following MIF stimulation, which was attenuated by 4-IPP, agmatine, or thiamine (**Fig. 4C**). Together, these results indicate that MIF promotes TM stiffness by driving fibrotic remodeling, ECM accumulation, and cytoskeletal reorganization, and that pharmacologic or immunometabolic inhibition of MIF can attenuate these pathological responses.

**Fig. 4:**
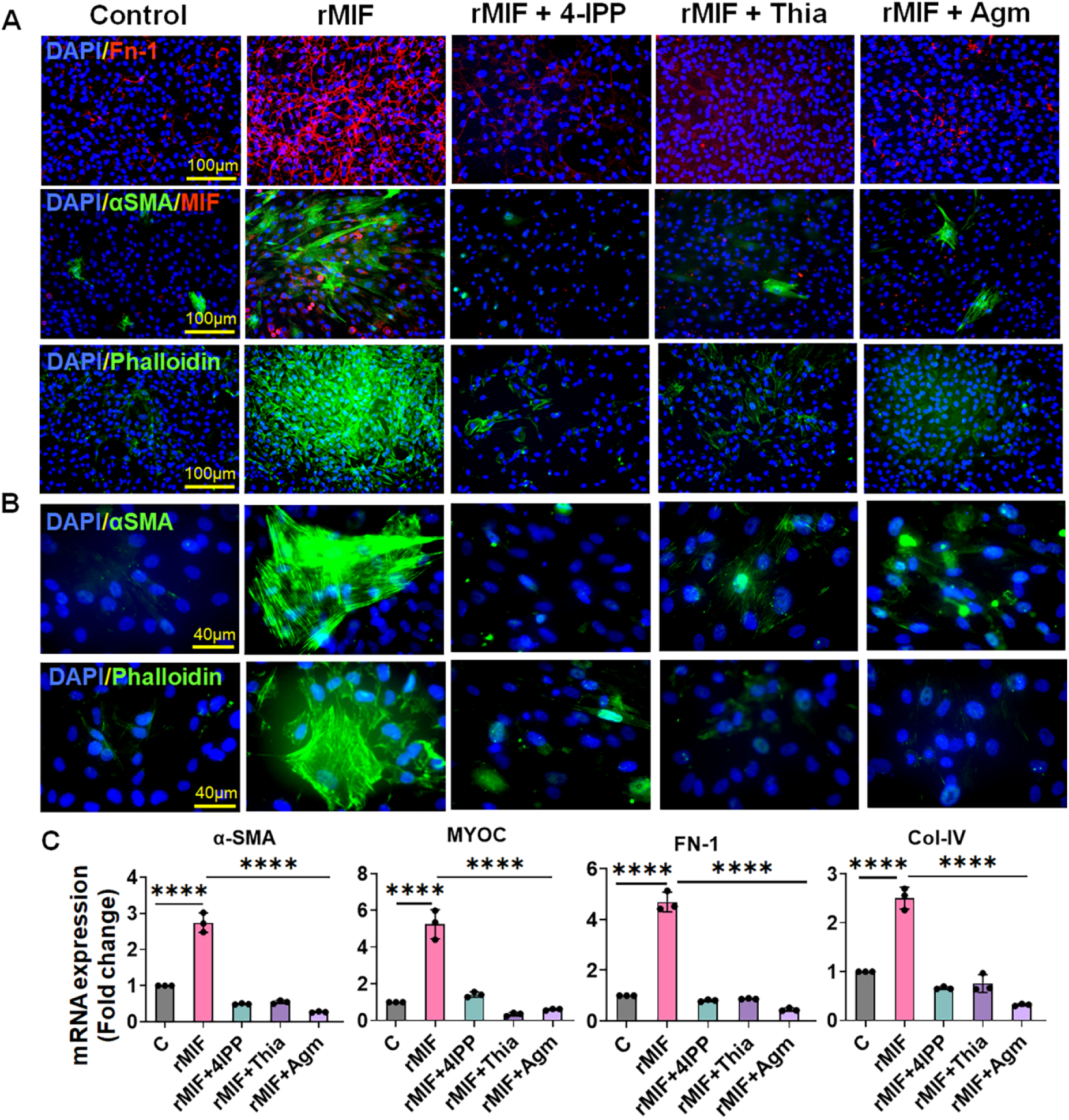
MIF promotes, while MIF inhibition attenuates, ECM deposition and cytoskeletal remodeling in HTMCs. Primary HTMCs (n=3) were challenged with rMIF (100ng/mL) in the presence or absence of the MIF inhibitor 4-IPP (100µM) or the metabolites agmatine (Agm) or thiamine (Thia) (100 ng/mL) for 24h. **(A-B)** Immunofluorescence analysis shows that rMIF increases ECM deposition, as indicated by increased FN-1 staining, and induces cytoskeletal alterations, evidenced by increased α-SMA and F-actin (phalloidin) staining. These pathological responses are markedly reduced by MIF inhibition with 4-IPP, Agm, or Thia. Scale bars: 100 µm (A), 40 µm (B). **(C)** qPCR analysis demonstrates that MIF significantly upregulates mRNA expression of fibrotic and ECM-related genes, including α-SMA, MYOC, FN-1, and COL-IV, whereas MIF inhibition attenuates these responses. Data represent mean ± SD. ****p<0.0001, One-way ANOVA with Dunnett’s multiple-comparison test.

### MIF activates RhoA/Rock-mediated MLC phosphorylation, while MIF inhibition suppresses these pathways

The RhoA/ROCK pathway is a key regulator of TM contractility, modulating cytoskeletal dynamics and ECM synthesis. Various pathological stressors, including inflammation, mechanical strain, and ECM accumulation, activate Rho-GTPase through guanine nucleotide exchange factors, leading to ROCK activation and subsequent MLC phosphorylation. Phosphorylated MLC enhances actomyosin cross-bridging, promoting cytoskeletal reorganization and TM cell contraction, thereby increasing aqueous humor outflow resistance and elevating IOP^28, 29^.

To determine whether MIF modulates this contractile pathway, we challenged primary HTMCs with rMIF in the presence or absence of the MIF inhibitor 4-IPP or the immunomodulatory metabolites agmatine and thiamine. We assessed the RhoA/ROCK activation and MLC phosphorylation by western blotting. Our results show that MIF exposure activated the RhoA/ROCK signaling and increased MLC phosphorylation (**Fig. 5A, B**). In contrast, inhibition of MIF with 4-IPP, agmatine, or thiamine markedly suppressed RhoA/ROCK activation and reduced MLC phosphorylation. To further validate these findings, myosin II activation was examined by immunofluorescence staining for phosphorylated MLC. To this end, our data show MIF-treated HTMCs increased pMLC staining, indicating enhanced myosin activity, whereas MIF inhibition substantially reduced pMLC levels (**Fig. 5C**). We used TGF-β2-treated cells as a positive control, which demonstrated comparable induction of pMLC activity. Collectively, these results demonstrate that MIF modulates TM contractility through activation of the RhoA/ROCK/MLC pathway. This contractile remodeling likely contributes to increased outflow resistance and elevated IOP, supporting MIF as a key pathological mediator of TM dysfunction in glaucoma.

**Fig. 5:**
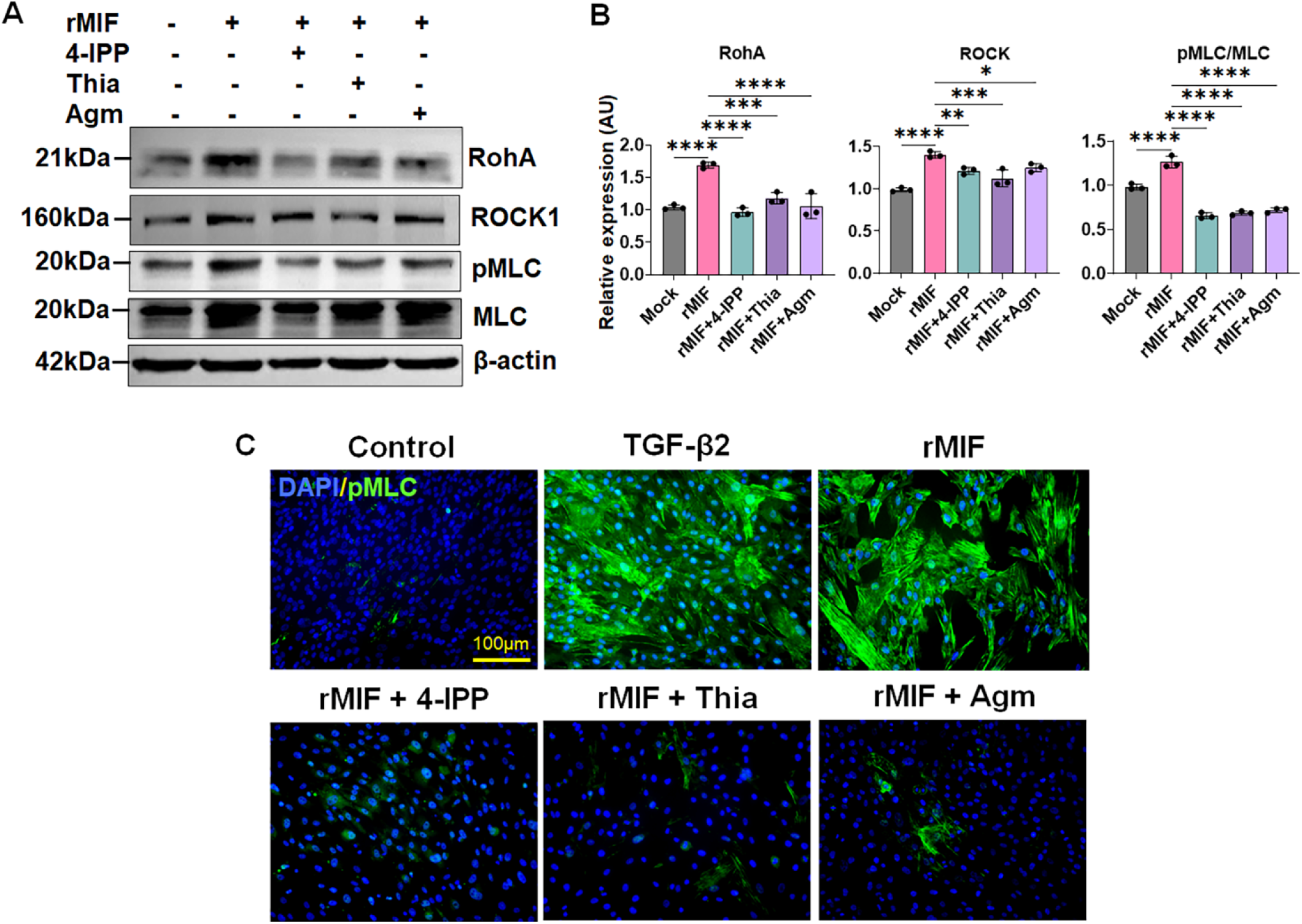
MIF activates the RhoA/ROCK pathways and induces MLC phosphorylation in HTMCs. Primary HTMCs (n=3) were challenged with rMIF (100 ng/mL) in the presence or absence of MIF inhibitor 4-IPP (100µM) or the metabolites Agm or Thia (100 ng/mL) for 24h. Cells treated with TGF-β2 (10ng/mL) served as a positive control. **(A-B)** Western blot analysis confirms activation of the RhoA/ROCK/pMLC signaling pathway in response to MIF treatment, which is significantly attenuated by MIF inhibition with 4-IPP, Agm, or Thia (**A**). Western blot bands were quantified by densitometric analysis using ImageJ and normalized to the housekeeping control β-actin, and protein levels are presented as relative expression (AU) (**B**). **(C)** Immunofluorescence analysis shows that MIF induces pMLC staining, comparable to that observed with TGF-β2 stimulation, which is significantly attenuated by MIF inhibition with 4-IPP, Agm, or Thia. Data represent mean ± SD. *p<0.05, **p<0.005, ***p<0.0005, and ****p<0.0001, One-way ANOVA with Dunnett’s multiple-comparison test.

### MIF inhibition restores Blimp-1 expression, suppresses inflammatory and cell death signaling, and enhances anti-inflammatory IL-10 response

Blimp-1 is a transcription factor that regulates the expression of the anti-inflammatory cytokine IL-10^22, 23^. Although MIF has been shown to suppress IL-10 production, whether MIF directly modulates Blimp-1/IL-10 has not previously been explored in any disease model. Our *in vivo* and *in vitro* findings demonstrated a consistent inverse relationship between MIF and Blimp-1, with elevated MIF levels accompanied by reduced Blimp-1. These observations led us to hypothesize that MIF negatively regulates the Blimp-1/IL-10 axis.

To test this, we challenged primary HTMCs with rMIF in the presence and absence of the MIF inhibitor 4-IPP and assessed the Blimp-1 expression by qPCR and immunoblotting. Surprisingly, MIF exposure significantly suppressed Blimp-1 expression, whereas pharmacologic inhibition of MIF with 4-IPP restored Blimp-1 levels at both the protein (**Fig. 6A**) and the transcript levels (**Fig. 6B**). Notably, treatment with immunomodulatory metabolites agmatine and thiamine similarly increased Blimp-1 expression, coinciding with suppression of MIF and CD74 (**Fig. 6A, B**). Given the established role of Blimp-1 in regulating IL-10, we next quantified IL-10 secretion by ELISA. Inhibition of MIF with 4-IPP, agmatine, or thiamine significantly increased IL-10 levels compared with rMIF-treated HTMCs (**Fig. 6C**), indicating restoration of anti-inflammatory signaling.

**Fig. 6:**
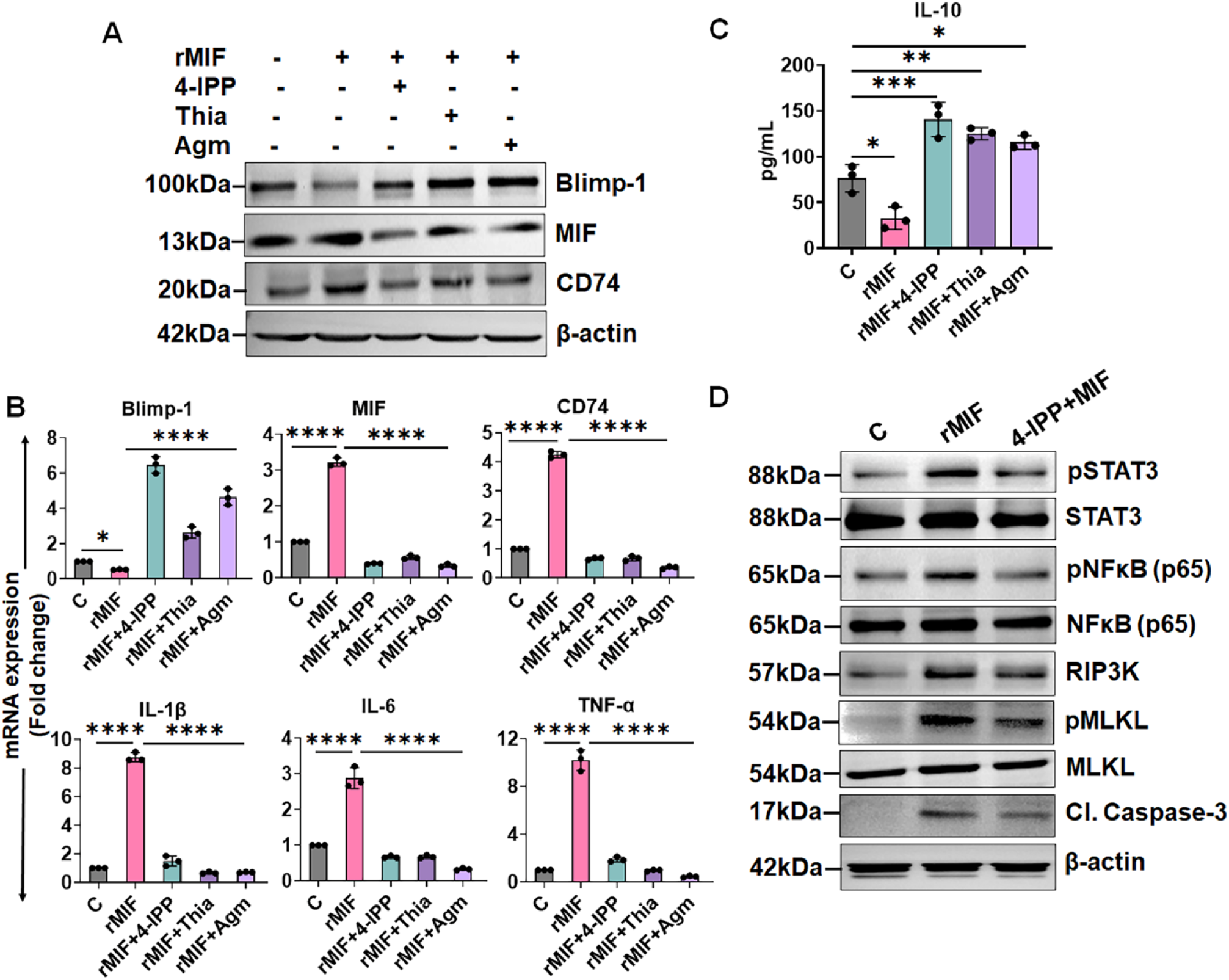
MIF inhibition restores Blimp-1/IL-10-mediated anti-inflammatory signaling and attenuates inflammatory and cell death responses in HTMCs. Primary HTMCs (n=3) were challenged with rMIF (100ng/mL) in the presence or absence of the MIF inhibitor 4-IPP (100µM) or the metabolites Agm or Thia (100 ng/mL) for 24h. **(A)** Western blot analysis shows that MIF suppresses Blimp-1 expression, whereas treatment with 4-IPP, Thia, or Agm restores Blimp-1 protein levels. **(B)** qPCR analysis reveals that MIF significantly induces mRNA expression of MIF, CD74, and pro-inflammatory cytokines (IL-1β, IL-6, TNF-α), while reducing Blimp-1 expression. Inhibition of MIF significantly restores Blimp-1 expression and suppresses MIF-CD74 signaling and pro-inflammatory cytokine expression. **(C)** ELISA analysis of treated and untreated TM culture supernatants demonstrates that MIF suppresses IL-10 secretion, which is restored following MIF inhibition. **(D)** Western blot analysis shows activation of inflammatory (STAT3, NFκB) and cell death pathways (RIP3K, MLKL, and cleaved caspase-3) following MIF treatment, which is markedly attenuated by MIF inhibition. Data represent mean ± SD. *p<0.05, and ****p<0.0001, One-way ANOVA with Dunnett’s multiple-comparison test.

We further examined the effect of MIF on inflammatory and cell death pathways. To test this, we challenged primary HTMCs with rMIF in the presence or absence of 4-IPP and measured the activation of inflammatory and cell death pathways by western blotting. Our data revealed that MIF significantly activated inflammatory (NFκB, STAT3) and cell death (MLKL, RIP3K, cleaved caspase 3) signaling pathways (**Fig. 6D**). In contrast, MIF inhibition markedly attenuated activation of these pathways. Consistent with these findings, our qPCR analysis also demonstrated that MIF exposure significantly increased expression of pro-inflammatory cytokines, including IL-1β, IL-6, and TNF-α (**Fig. 6B**). Inhibition of MIF with 4-IPP, agmatine, or thiamine significantly suppressed the transcription of these mediators.

Collectively, these results demonstrate that MIF suppresses the protective Blimp-1/IL-10 axis while promoting inflammatory and cell death signaling in TM cells. Pharmacologic or immunometabolic inhibition of MIF restores Blimp-1-mediated anti-inflammatory responses and attenuates pro-inflammatory and cell death pathways, supporting a central role for MIF in driving TM dysfunction.

### MIF inhibition reduces IOP and suppresses profibrotic and inflammatory responses in OHT Mice

Recently, we purified and characterized extracellular vesicles (EVs) derived from HTMCs and demonstrated their therapeutic potential for ocular drug delivery in a mouse model^27^. Here, to further assess the effect of MIF inhibition on IOP elevation and TM pathology *in vivo*, we treated LV-TGF-β2 OHT mice topically with EVs loaded with agmatine (EV-Agm) once daily for 9 weeks. IOP was recorded weekly in both treated and untreated groups. Our results show that EV-agmatine treatment significantly reduced IOP compared with LV-TGF-β2-injected, untreated controls (**Fig. 7A**). At 9 weeks post-treatment, TM/AS tissue were collected and analyzed by qPCR to assess for mRNA levels of profibrotic markers (α*-Sma, Fn-1, Myoc*), inflammatory mediators (*Il-1*β*, Il-6*), and *Mif-Cd74*/*Blimp-1* expression. Our results show that topical EV-Agm treatment significantly decreased *Mif* and *Cd74* expression, attenuated LV-TGF-β2-induced profibrotic and inflammatory responses, and restored *Blimp-1* expression (**Fig. 7B**). These findings indicate that MIF inhibition effectively mitigates TM dysfunction *in vivo*. Collectively, our results demonstrate that targeting MIF reduces IOP and alleviates TM pathology in experimental OHT, supporting the MIF-CD74/Blimp-1 axis as a potential therapeutic target in glaucoma.

**Fig. 7:**
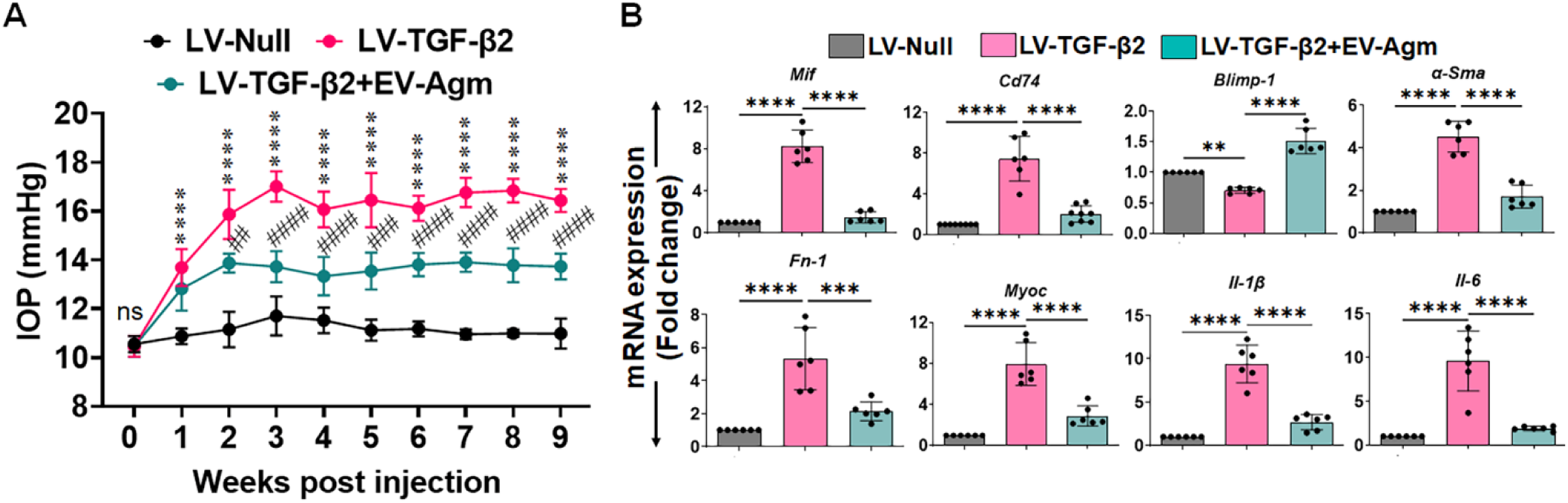
Topical EV-loaded agmatine reduces IOP and suppresses MIF-driven inflammatory and cytoskeletal/ECM responses in LV-TGF-β2-induced OHT. OHT was induced in WT C57BL/6 mice (n=12) by intravitreal injection of LV-TGF-β2. LV-null-injected mcie served as controls. Following OHT induction, extracellular vesicle-loaded agmatine (EV-Agm; 0.1 µg/eye) was applied topically once daily. **(A)** Weekly IOP measurements over 9 weeks show that EV-Agm treatment significantly reduces IOP compared with untreated LV-TGF-β2-injected mice. **(B)** qPCR analysis of anterior segment tissue demonstrates that EV-Agm significantly reduces expression of *Mif, Cd74,* pro-inflammatory cytokines (*Il-1*β*, Il-6*), and cytoskeletal/ECM markers (α*-Sma, Myoc, Fn-1*), while restoring *Blimp-1* expression compared with untreated LV-TGF-β2-injected mice. *p<0.05, **p<0.005, ###,***p<0.0005, and ####,****p<0.0001. Two-way ANOVA with Tukey’s multiple comparison (A), one-way ANOVA with Tukey’s multiple comparison (B). In panel A, * denotes LV-null vs. LV-TGF-β2, and # denotes LV-TGF-β2 vs. LV-TGF-β2+EV-Agm comparisons.

## Discussion

Elevated IOP resulting from TM dysfunction remains the primary modifiable risk factor for POAG. TM pathology is often characterized by tissue stiffening, ECM accumulation, cytoskeletal remodeling, and chronic low-grade inflammation, all of which contribute to increased aqueous humor outflow resistance and sustained IOP elevation^2, 30, 31^. Although current therapies effectively lower IOP, they do not directly address the underlying molecular drivers of TM dysfunction. Emerging evidence implicates immune and inflammatory mechanisms as critical contributors to TM pathology^32, 33^; however, the specific roles of innate inflammatory signaling pathways remain incompletely defined. In the present study, we identify MIF and its receptor CD74 as central mediators integrating fibrotic, biomechanical, and inflammatory processes in glaucomatous TM dysfunction. Using two complementary OHT mouse models and mechanistic studies in primary human TM cells *in vitro*, we demonstrate that glaucomatous stress induces MIF-CD74 signaling while concurrently suppressing the immunoregulatory transcription factor Blimp-1. Mechanistically, MIF promotes fibrotic remodeling, enhances RhoA/ROCK/pMLC-dependent TM contractility, and amplifies inflammatory and cell death signaling. Collectively, these findings identify MIF as a unifying upstream regulator of TM pathology.

MIF is a pleiotropic pro-inflammatory cytokine that signals primarily through CD74 and activates NFκB, MAPK, and STAT pathways in multiple inflammatory and autoimmune diseases^11, 19, 34^. Ocular expression of MIF in the TM was first reported more than two decades ago, showing that it modulates the Th1 cytokine response without increasing IL-10 secretion, suggesting a pro-inflammatory role in anterior segment tissues^13^. More recent studies have further highlighted the importance of the MIF-CD74 signaling pathway in ocular inflammatory diseases^17, 18, 35, 36^. However, the contribution of MIF to glaucomatous TM dysfunction has remained undefined. In this study, using mutant myocilin-induced OHT (Tg.CreMYOC^Y437H^)^25^ and lentiviral TGF-β2-induced OHT^37^, we demonstrate for the first time robust upregulation of MIF and CD74 in TM/AS tissues, coincident with sustained IOP elevation and induction of fibrotic markers. These findings suggest that MIF activation is not merely a secondary inflammatory response but may actively contribute to TM pathology.

TGF-β2 is widely recognized as a central profibrotic mediator in glaucoma. Elevated aqueous humor levels of TGF-β2 have been consistently reported in POAG patients, and its overexpression has been shown to induce ECM accumulation and OHT^26, 37–39^. Our data demonstrate that LV-TGF-β2-induced OHT is accompanied by marked activation of MIF-CD74 signaling and concurrent suppression of Blimp-1, suggesting that MIF may function downstream of, or in parallel with, TGF-β2 signaling to amplify profibrotic remodeling. Notably, mutant myocilin-driven OHT, which models proteotoxic stress-induced TM dysfunction, also triggered MIF upregulation, indicating that MIF activation represents a convergent response to distinct glaucomatous stressors. Consistent with these *in vivo* observations, our *in vitro* studies showed that ocular hypertensive agents, such as TGF-β2 and a pro-inflammatory milieu, induce MIF expression in primary HTMCs, supporting a stress-responsive role for MIF in TM under glaucomatous conditions. Our findings corroborate earlier reports demonstrating that MIF is expressed and secreted by human TM cells^13^ and is upregulated upon Cathepsin K depletion^40^, a protease critical for ECM degradation, highlighting its potential role as a key mediator linking profibrotic and inflammatory pathways in glaucoma.

Fibrotic ECM accumulation and increased TM stiffness are central determinants of elevated outflow resistance in POAG^30, 31, 41^. Excessive fibronectin deposition and induction of α-SMA promote a myofibroblast-like phenotype associated with increased tissue rigidity. Our findings demonstrate that MIF directly induces α-SMA expression, fibronectin deposition, and F-actin cytoskeletal reorganization in HTMCs, establishing MIF as a profibrotic mediator in TM cells. Importantly, pharmacologic inhibition of MIF with 4-IPP or metabolic modulation with agmatine and thiamine, metabolites we recently identified as altered in glaucoma patients^27^, significantly attenuated these pathogenic responses. Notably, our study establishes, for the first time, that agmatine and thiamine suppress MIF activity and its associated pro-inflammatory and profibrotic effects. Together, these results extend the current paradigm of TM fibrosis by identifying MIF as a non-canonical amplifier of ECM and cytoskeletal remodeling that complements established TGFβ-dependent pathways.

In addition to ECM remodeling, TM contractility plays a critical role in regulating aqueous humor outflow. The RhoA/ROCK pathway is a key modulator of TM cytoskeletal dynamics; its activation promotes MLC phosphorylation, enhances actomyosin cross-bridging, and increases cellular contractility^29, 42^. Pharmacologic inhibition of ROCK lowers IOP by relaxing TM cells and improving outflow facility, forming the basis for clinically approved ROCK inhibitors^42–44^. Our results demonstrate that MIF robustly activates RhoA/ROCK signaling and increases MLC phosphorylation in HTMCs, whereas MIF inhibition suppresses these contractile responses. These findings suggest that MIF integrates inflammatory signaling with biomechanical dysregulation, acting upstream of RhoA activation and linking cytokine-driven stress to increased TM stiffness and outflow resistance.

A particularly novel aspect of our study is the identification of the Blimp-1/IL-10 axis as a downstream regulatory target of MIF signaling in TM cells. Blimp-1 is a transcriptional regulator essential for IL-10 expression and maintenance of anti-inflammatory homeostasis. Although MIF has been reported to suppress IL-10 production in TM and other immune cells^12, 13, 34^, direct regulation of Blimp-1 by MIF has not previously been demonstrated in any disease model. Our findings reveal a consistent inverse relationship between MIF and Blimp-1 expression in both OHT models and in HTMCs exposed to glaucomatous stress. MIF suppressed Blimp-1 expression and reduced IL-10 secretion, whereas pharmacologic inhibition of MIF restored Blimp-1 levels and enhanced IL-10 production. Given that pro-inflammatory cytokines such as IL-1β, IL-6, and TNF-α are elevated in glaucomatous TM and aqueous humor^45–47^, our findings suggest that MIF not only promotes inflammatory signaling but also suppresses endogenous anti-inflammatory counter-regulation through inhibition of the Blimp-1/IL-10 axis. Consistent with this interpretation, MIF enhanced activation of NFκB and STAT3 pathways and increased markers of necroptotic and apoptotic signaling, including RIPK3, MLKL, and cleaved caspase-3, supporting a role in TM cell death and degeneration.

Taken together, our findings support a unified model in which glaucomatous stress, whether genetic (mutant myocilin) or cytokine-driven (TGF-β2), induces activation of the MIF-CD74 pathways in the TM. MIF activation promotes ECM accumulation and myofibroblast-like transformation, enhances RhoA/ROCK-mediated contractility, and suppresses anti-inflammatory Blimp-1/IL-10 axis while simultaneously activating NFκB/STAT3 and cell death pathways. These interconnected mechanisms increase tissue stiffness, elevate outflow resistance, and perpetuate chronic inflammatory signaling, collectively driving IOP elevation and progressive TM dysfunction. Importantly, pharmacologic or immunometabolic inhibition of MIF reduced IOP in OHT mice and attenuated both profibrotic and inflammatory responses within the TM, highlighting the therapeutic reversibility of this pathway. While our study provides compelling mechanistic insights into the role of MIF in TM dysfunction, additional studies are needed to establish causality. Future studies using genetic MIF gain- and loss-of-function approaches will help clarify its direct contribution to TM dysfunction and outflow resistance. It will also be important to determine whether MIF signaling operates independently of TGF-β2 or functions cooperatively within established profibrotic networks, including SMAD signaling and integrin-mediated mechanotransduction. Furthermore, the potential involvement of alternative MIF receptors and receptor-independent biological actions in TM biology also warrants further exploration.

In summary, our study identifies MIF as a central upstream mediator integrating fibrotic remodeling, biomechanical dysregulation, and inflammatory signaling in the TM. By coupling ECM accumulation and RhoA/ROCK-dependent contractility with suppression of the Blimp-1/IL-10 anti-inflammatory axis, MIF orchestrates a coordinated pathogenic program that promotes TM dysfunction and sustained IOP elevation. Targeting MIF signaling may therefore represent a promising therapeutic strategy aimed not only at lowering IOP but also at restoring TM homeostasis and modifying the underlying molecular mechanisms that drive glaucoma progression.

## Materials & Methods

### Reagents

Recombinant human proteins: TNF-α (#300-01A-50UG), IL-1β (#200-01B-50UG), IL-6 (#200-06-50UG), IL-17A (#200-17-50UG), and MIF (#300-69-25UG) were obtained from PeproTech (ThermoFisher Scientific, Rockford, IL). Recombinant human TGF-β2 (302-B2-010/CF) protein was purchased from R&D Systems (Minneapolis, MN). The MIF inhibitor, 4-iodo-6-phenylpyrimidine (4-IPP; #31770), agmatine (#23513), and thiamine (#25656) were obtained from Cayman Chemicals (Ann Arbor, MI). Primary antibodies used in this study were purchased from the following sources: β-actin (Millipore Sigma, #A2228), pNFκB (#3033), NFκB (#6956), pSTAT3 (#9145), STAT3 (#9139), RIP3K (#10188), pMLKL (#18640), MLKL (#14993), RhoA (#2117), ROCK1 (#4035), pMLC (#3674), MLC (#3672), MIF (#75038), CD74 (#77274), Blimp-1 (#9115) (Cell Signaling Technology, Danvers, MA). Recombinant adenoviral vectors HAd5-Cre and HAd5-empty were obtained from the Viral Core Facility at the University of Iowa. Lentiviral vectors encoding the active hTGF-β2 (C226S/C228S) under the control of CMV promoter (LV-TGF-β2) and corresponding empty vector control (LV-null), were purchased from VectorBuilder (Chicago, IL).

### Cell Culture and Treatment Conditions

Validated normal primary human trabecular meshwork cells (HTMCs)^48–50^ were cultured in Dulbecco’s modified Eagle’s medium (DMEM) supplemented with 10% fetal bovine serum (FBS) and 1X penicillin-streptomycin (ThermoFisher Scientific, Rockford, IL) at 37°C in a humidified incubator with 5% CO_2_. Cells were treated with TGF-β2 (10 ng/mL), rMIF (100 ng/mL), or a proinflammatory cytokine cocktail (TNF-α, IL-1β, IL-6, and IL-17A; 100 ng/mL) for 24h. For MIF inhibition studies, cells were pretreated with the specific irreversible MIF inhibitor 4-IPP (100μM), or agmatine or thiamine (100ng/mL) for 2h prior to MIF stimulation.

### Animals and Ethics Statement

Wild-type (WT) C57BL/6 mice (strain #:000664) were obtained from Jackson Laboratory (Bar Harbor, ME). Tg.CreMYOC^Y437H^ mice^25^ were generously provided by Dr. Gualb Zode, Professor, University of California, Irvine. Mice were bred in-house in a germ-free University of Missouri (MU) Office of Animal Resources (OAR) facility and genotyped. Both male and female mice aged 6-12 weeks were used in this study. Animals were housed in a controlled-access, AAALAC-accredited OAR facility under a 12:12h light/dark cycle with ad libitum access to rodent chow (LabDiet Pico Laboratory, St. Louis, MO) and water. All procedures were conducted in accordance with the Association for Research in Vision and Ophthalmology (ARVO) Statement for the Use of Animals in Ophthalmic and Vision Research. Experimental and biohazard protocols were approved by the MU Animal Care and Use Committee (ACUC) and Institutional Biosafety Committee (IBC).

### OHT induction in mice

Heterozygous Tg.CreMYOC^Y437H^ mice were anesthetized and intravitreally injected with HAd5-cre (2 x 10^7^ pfu/eye) to induce expression of the MYOC^Y437H^ mutation^25^. Mice injected with an equivalent dose of HAd5-empty vector served as controls. For TGF-β2-induced OHT, WT C57BL/6 mice were intravitreally injected with a lentiviral vector encoding constitutively active human TGF-β2 (C226S/C228S) under the CMV promoter (LV-TGF-β2; 2 × 10 TU/eye in a 2 µL bolus)^37^. Mice injected with an equivalent amount of empty lentiviral vector (LV-null) served as controls.

### IOP Measurement

Following OHT induction, IOP was recorded weekly using TonoLab rebound tonometer (Colonial Medical Supply, Espoo, Finland), as previously described^48, 50^. IOP measurements were performed on conscious mice during the daytime (10:00 AM-12:00 PM) to minimize diurnal variation. To ensure consistency, all recordings were obtained at the same time of day. Baseline IOP was recorded prior to intravitreal injections, and subsequent measurements were performed weekly thereafter. For each eye, six consecutive measurements were obtained, with each measurement representing the average of six individual rebound readings as determined by the tonometer. All IOP assessments were performed in a masked manner.

### Western Blot

Total proteins were extracted from HTMCs or anterior segment tissue or corneoscleral TM rims using RIPA lysis buffer (ThermoFisher Scientific, Rockford, IL). Protein concentrations were determined using a bicinchoninic acid (BCA) assay (ThermoFisher Scientific, Rockford, IL). Equal amounts of protein (30 μg) were resolved on 8-14% SDS-polyacrylamide gels and transferred onto polyvinylidene difluoride (PVDF) membranes (ThermoFisher Scientific, Rockford, IL). Membranes were blocked with 5% nonfat milk in TBST at room temperature (RT) for 1 h and incubated overnight at 4°C with respective primary antibodies (1:1000; β-actin, 1:5000). The following day, membranes were washed three times with TBST and incubated with corresponding horseradish peroxidase-conjugated secondary antibodies (1:2000; β-actin, 1:10,000) for 2h at RT. Proteins bands were detected using SuperSignal West Femto Maximum Sensitivity substrate and imaged with the iBright FL1500 gel documentation system (ThermoFisher Scientific, Rockford, IL). Densitometric analysis was performed using ImageJ software (NIH)^51^.

### Immunofluorescence Staining

Primary HTMCs were cultured on Nunc four-well chamber slides (Fisher Scientific, Rochester, NY) and treated with rMIF (100 ng/mL), TGF-β2 (10 ng/mL), and/or the MIF inhibitor 4-IPP to assess α-SMA and pMLC expression. For inhibition studies, cells were pretreated with 100 μM 4-IPP for 2 h prior to 24 h MIF induction. For both *in vitro* and ocular tissue staining, samples were fixed in 4% paraformaldehyde in PBS at 4°C, washed three times with PBS, and blocked/permeabilized for 1h at RT using either 1% (w/v) BSA with 0.4% Triton X-100 (HTMC slides) or 10% (v/v) goat serum with 0.4% Triton X-100 (ocular tissue sections). Samples were then incubated overnight at 4°C with respective primary antibodies (1:100). Following four PBS washes, cells and tissue sections were incubated with anti-mouse/rabbit Alexa Fluor 488- or 594-conjugated secondary antibodies (1:200) for 1 h at RT. After four final washes, coverslips were mounted with Vectashield antifade mounting medium containing DAPI (Vector Laboratories, Burlingame, CA) and imaged using a Keyence microscope (Keyence, Itasca, IL).

### RNA Extraction and Quantitative RT-PCR

Total RNA was extracted from treated and untreated cells or tissue samples using TRIzol reagent, following the manufacturer’s instructions (ThermoFisher Scientific, Rockford, IL). cDNA was synthesized from 1 µg of total RNA using the Maxima first-strand cDNA Synthesis Kit, per the manufacturer’s instructions (ThermoFisher Scientific, Rockford, IL). qPCR was performed using human or mouse gene-specific primers (**Table 1**) on a QuantStudio 3 Real-Time PCR system (ThermoFisher Scientific, Rockford, IL). Relative gene expression levels were normalized to the constitutive 18S rRNA and analyzed using the ΔΔC_T_ method^52^, and results are expressed as fold change relative to controls.

**Table 1:**
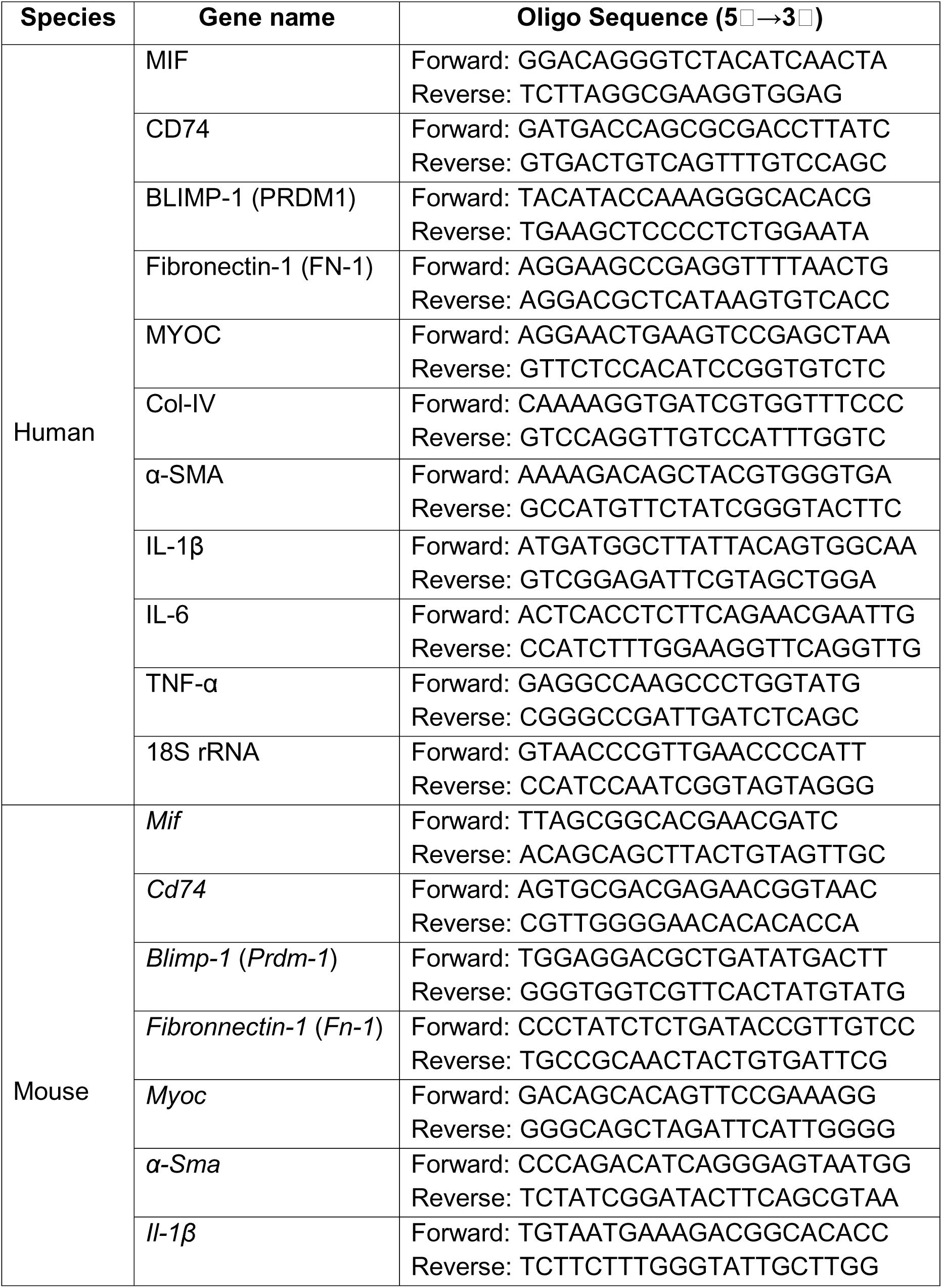
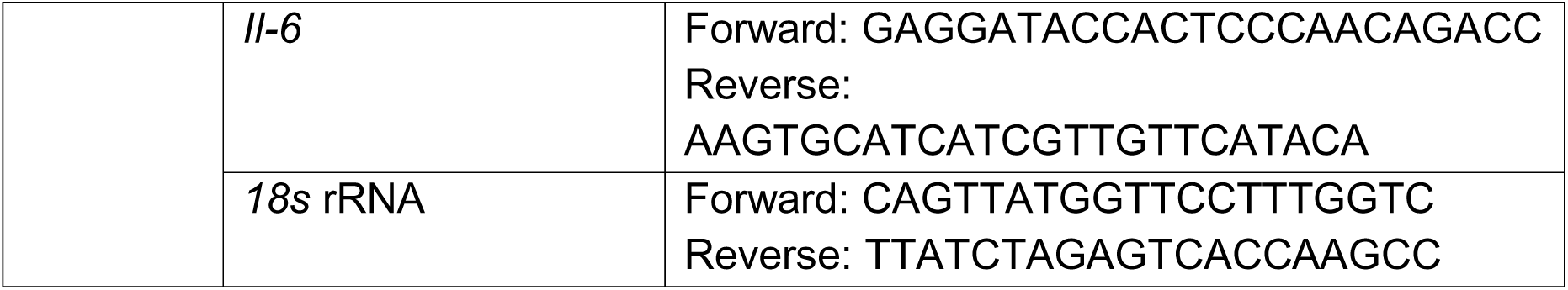
List of primers and their gene sequences used for qPCR.

### Statistical Analyses

All statistical analyses were performed using GraphPad Prism 10.1.2 (GraphPad Software, San Diego, CA). Comparisons between two groups were made using an unpaired t-test. For comparisons of three or more groups, either one-way or two-way ANOVA was applied with appropriate post hoc tests. All data are presented as the mean ± standard deviation (SD) from at least three independent experiments unless otherwise specified. A p-value of <0.05 was considered statistically significant.

## Acknowledgements

This study is supported in part by the National Institute of Health (NIH)/ National Eye Institutes (NEI) grant R01EY032495 and research start-up funds from the University of Missouri School of Medicine to PKS and R01EY026177 to GZ. The funders had no role in study design, data collection and analysis, decision to publish, or preparation of the manuscript. We are grateful to Dr. Akhil Srivastava, Assistant Professor at the University of Missouri, and his team for their assistance with extracellular vesicle preparation and agmatine loading. The authors also thank the staff of the MU Office of Animal Resources for their excellent care and management of the animal colonies.

## Conflict of Interest

The authors declare no conflict of interest.

## Data Availability

All relevant data supporting the findings of this study are included in the manuscript and figures.

## Author Contributions

PKS conceived the study, designed and supervised the experiments, and secured funding for the study. MM, LKK, and PK performed experiments and analyzed the data. MM and PKS wrote the manuscript. GZ provided Tg.CreMYOC^Y437H^ mice, primary HTMCs, and critical intellectual input. All authors reviewed, edited, and approved the final manuscript.

## Notes

### Competing Interest Statement

The authors have declared no competing interest.

